# Dietary omega-3 polyunsaturated fatty acids interact with select gut microbes to produce bioactive metabolites found in infant gut microbiomes

**DOI:** 10.64898/2026.10.10.756005

**Authors:** Janine M.C. Comrie, Xiaoqing Tan, Henry H. Le, Min-Ting Lee, Annett Richter, Sharon C. Thompson, Kevin R. Besler, Elizabeth L. Johnson

**Affiliations:** Division of Nutritional Sciences, Cornell University, Ithaca, NY; Department of Microbiology, Cornell University, Ithaca, NY; Howard Hughes Medical Institute, Cornell University, Ithaca, NY

## Abstract

Gut microbes metabolize nutrients in infant diets, producing compounds that play roles in host health. Long-chain polyunsaturated fatty acids (LCPUFAs) are an important component of infant diets, supporting neurological development and immune regulation. Emerging evidence demonstrates that gut microbes, alongside the host, participate in the metabolism of dietary lipids, highlighting an underexplored avenue for LCPUFA biology. In this work, alkyne-tagged eicosapentaenoic acid (EPA^Alk^) was used to investigate interactions between LCPUFAs and the gut microbiota. Flow cytometry and comparative metabolomics identified gut microbial EPA interactors in the ileum of mice. Further comparative metabolomics analysis identified the conversion of EPA to *N*-acyl amino acids, including attachment to phenylalanine (EP-Phe) and leucine (EP-Leu). Targeted analysis of *ex vivo* stool cultures determined that EP-Phe and EP-Leu are produced by gut bacteria. A survey of lipid-relevant infant gut bacteria determined various flow cytometry-confirmed EPA-microbe interactions and identified *Enterococcus faecalis, Bacteroides thetaiotaomicron,* and *Clostridium bifermentans* as competent producers of EP-Phe or EP-Leu. Thus, both novel metabolites from diet-derived inputs and microbial producers of these metabolites were identified. Together, this highlights important microbiome-centric consequences of LCPUFA consumption and emphasizes that the gut microbiome may be an important consideration in targeted supplementation strategies.

---

Dietary long-chain polyunsaturated fatty acids (LCPUFAs) support diverse biological functions, and inadequate or imbalanced intake may affect human health. Notably, LCPUFAs contribute to infant development, particularly by shaping metabolism^1^, neurodevelopment^2,3^ and immune function^4-7^. After birth, infants can acquire LCPUFAs through human milk^8^, including docosahexaenoic acid (DHA) and arachidonic acid (ARA), which accumulate rapidly in the developing brain during early infancy^3,9-11^. The importance of DHA and ARA to infant biology has led to the inclusion of DHA-rich algal oil and ARA-rich fungal oil in U.S. infant formulas since the early 2000s^12,13^ despite the only regulatory requirement for LCPUFAs being the inclusion of linoleic acid in U.S. formulations^14^. Supplementation studies report some beneficial developmental outcomes with LCPUFA supplementation, particularly in preterm infants, including improved cognitive development and reduced gut pathogen burden^15,16^. Although supplementation studies highlight potential developmental influences of LCPUFAs, the role of the gut microbiome in mediating these effects remains understudied and may contribute to the variability in outcomes observed across LCPUFA supplementation studies^17^.

Work in mouse model systems, microbial culturing, and human supplementation trials provides mechanistic support for the importance of dietary LCPUFAs in influencing host biology through the microbiome. Upon consumption, LCPUFAs reach the distal small intestine, where they interact with commensal microbes^18^. LCPUFA consumption for both infants and adults has been shown to alter microbiome composition^16,19,20^. The most extensively studied microbial interactions with LCPUFAs involve linoleic acid (LA) and α-linolenic acid (ALA) which are converted to hydroxy, oxo-, and conjugated fatty acids^21-24^. Products of LA have been detected in the small intestine of mice, where these metabolites are absorbed^21^. Studies supplementing mice with an LA-derived hydroxy-fatty acid show reduced inflammation in various tissues and restoration of gut integrity^25,26^. Additionally, oxo-fatty acids from both LA and ALA have been shown to induce PPARγ signaling^23,27^. Signaling events driven by ALA-derivatives were shown to reduce inflammatory activity in both skin and adipose tissue in mice^27^. Finally, LA-derived oxo-fatty acids have been shown to influence cell differentiation and glucose uptake in adipocytes^23^. The bioactivity of LA- and ALA-derived metabolites in these model systems provides compelling evidence that gut microbial LCPUFA metabolism can exert health-modifying effects on host physiology.

Given microbial metabolism can transform dietary components into bioactive compounds that modulate host physiology, identifying novel dietary LCPUFA-microbe interactions can provide critical information on the efficacy of supplementation efforts. Identifying specific microbes that are responsive to dietary LCPUFA exposure and the microbial metabolites created through microbial uptake of diet-derived LCPUFAs can clarify infant nutritional and microbial supplementation recommendations. Efforts to identify dietary LCPUFA-dependent microbes could provide important resources for increasing the efficacy of dietary supplement-based health interventions. This is especially critical in the neonatology intensive care unit (NICU) where antibiotic use, other medicines, and care practices can lead to varied microbiome compositions in infants and could ultimately contribute to the varied effects of dietary LCPUFA consumption on infant health outcomes^28^. To investigate the potential for interindividual variability in gut microbiota LCPUFA exposure, we sought to characterize the LCPUFA landscape of infant feeding modes and observed a significant difference in the available eicosapentaenoic acid (EPA) in human milk as compared to infant formula. We then focused on investigating dietary EPA-microbiota interactions by using a chemical biology-based workflow to identify the production of novel microbial EPA-derived metabolites.

## RESULTS

### LCPUFA levels in human milk and select infant formulas

Dietary LCPUFA exposure can vary between infant feeding modes^29,30^. To quantify dietary LCPUFAs entering the infant gut across feeding modes, we determined the absolute concentration of LCPUFAs in human milk across multiple mothers and in various infant formulas. Free fatty acid LCPUFA concentrations and total LCPUFA concentrations were determined for 8 infant formulas and for mature human milk from 16 mothers using high-resolution liquid chromatography-mass spectrometry (LC-MS) (Figure 1A-B). Because many fatty acids, including LCPUFAs, are sequestered in complex lipids, we deployed a hydrolytic approach to enable quantification of esterified LCPUFAs (Figure 1B). The abundances of free fatty acid EPA, DHA, ARA, and LA were substantially higher in human milk compared to infant formulas (Figure 1B). After lipid hydrolysis, total LCPUFA concentrations in infant formula were similar to or higher than those observed in human milk (Figure 1B). Given the staged development of lipid digestion during early infancy^31^, the availability of free fatty acids is a notable difference between these dietary sources. Furthermore, we note that different forms of LCPUFAs across dietary modes may influence their availability to the gut microbiota.

**Figure 1.**
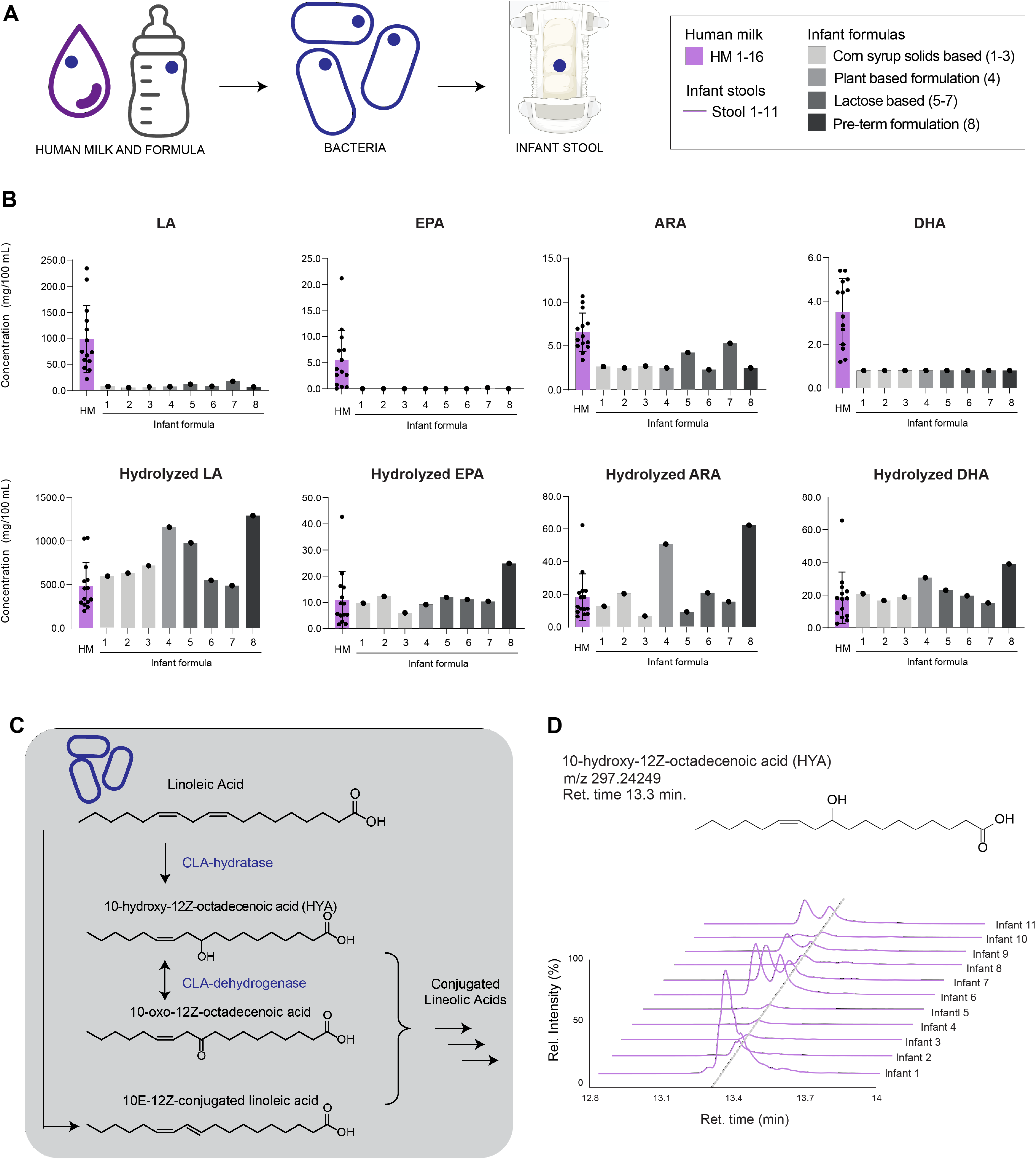
LCPUFA levels in human milk versus infant formula and detection of a microbe-associated LCPUFA derivative in infant stool.

A) Scheme for nutrients in infant diet transferring to gut microbiota and then to stool. Dark blue circles indicate nutrient/nutrient-derived microbial metabolite of interest. (B) Concentrations of free, non-esterified EPA, DHA, ARA, and LA were measured in 16 human milk and in 8 different infant formulas using high-resolution LC-MS. Concentrations of total EPA, DHA, ARA, and LA, freed through lipid hydrolysis, were measured in the same human milk samples and infant formulas. (C) Pathway for the conversion of LA to 10-hydroxy-12(Z)-octadecenoic acid (HYA) by *Lactobacillus* species. (D) Extracted ion chromatograms (EICs) depicting HYA detection in human infant stool using high-resolution LC-MS.

### Infant gut microbial capacity to metabolize LCPUFAs cooccurs with abundance of known LCPUFA-dependent microbial metabolites

To determine if infant microbial communities metabolize LCPUFAs, we focused on the known transformation of LA and ALA to hydroxy-and conjugated fatty acids by gut-associated *Lactobacillus* and *Bifidobacterium* spp. (Figure 1C)^21-24^. In these bacterial systems, LA is converted to 10-hydroxy-12(Z)-octadecenoic acid (HYA). Indeed, in human infant stool, we detected HYA, suggesting this gut bacterial metabolic activity is present in infants (Figure 1D). Additionally, shotgun metagenomic sequencing revealed putative HYA-producing bacterial species in these stool samples, further suggesting LCPUFA-gut microbe interactions occurred (Figure S1). While additional studies in larger cohorts are required to further elaborate the HYA, *Lactobacillus,* and *Bifidobacterium* association, these results motivated more detailed analysis of the metabolic interactions among the gut microbiome, diet-derived LCPUFAs, and human infants. Particularly, we chose to focus on EPA because it is an understudied LCPUFA in infant nutrition. Findings regarding EPA have direct applications to DHA- and ARA-derived microbial metabolites, making it an attractive candidate for further analysis. Thus, we chose to focus on further exploration of potential interactions of dietary EPA with the gut microbiome.

### Gut bacterial EPA utilization and EP amino acids detected *in vivo*

To reveal EPA-gut microbiome interactions, we leveraged our recently developed workflow that allows for detailed tracing of nutrient-microbiota interactions in host systems. This methodology uses techniques in bioorthogonal chemistry and comparative metabolomics to understand both the microbes that take up dietary inputs and the microbial metabolites that result from nutrient-microbiota interactions. We used an alkyne-modified form of EPA that has an alkyne (C≡C) at the omega position (EPA^Alk^) to trace the fate of dietary EPA within the gut microbial community. To determine novel diet-derived EPA interactions with the gut microbiome, we introduced EPA^Alk^ to mice via oral gavage once daily for 3 consecutive days (Figure 2A). Microbes that utilized diet-derived EPA^Alk^ were detected using click chemistry to covalently link a fluorophore (Alexa Fluor 647-Azide, red) to alkyne-containing metabolites consumed by gut microbes. Bacterial cells isolated from the intestinal contents of EPA^Alk^-treated mice displayed red fluorescence as detected by microscopy (Figure 2B) and flow cytometry (Figure 2C), highlighting a subset of bacteria that engaged diet-derived EPA^Alk^.

**Figure 2.**
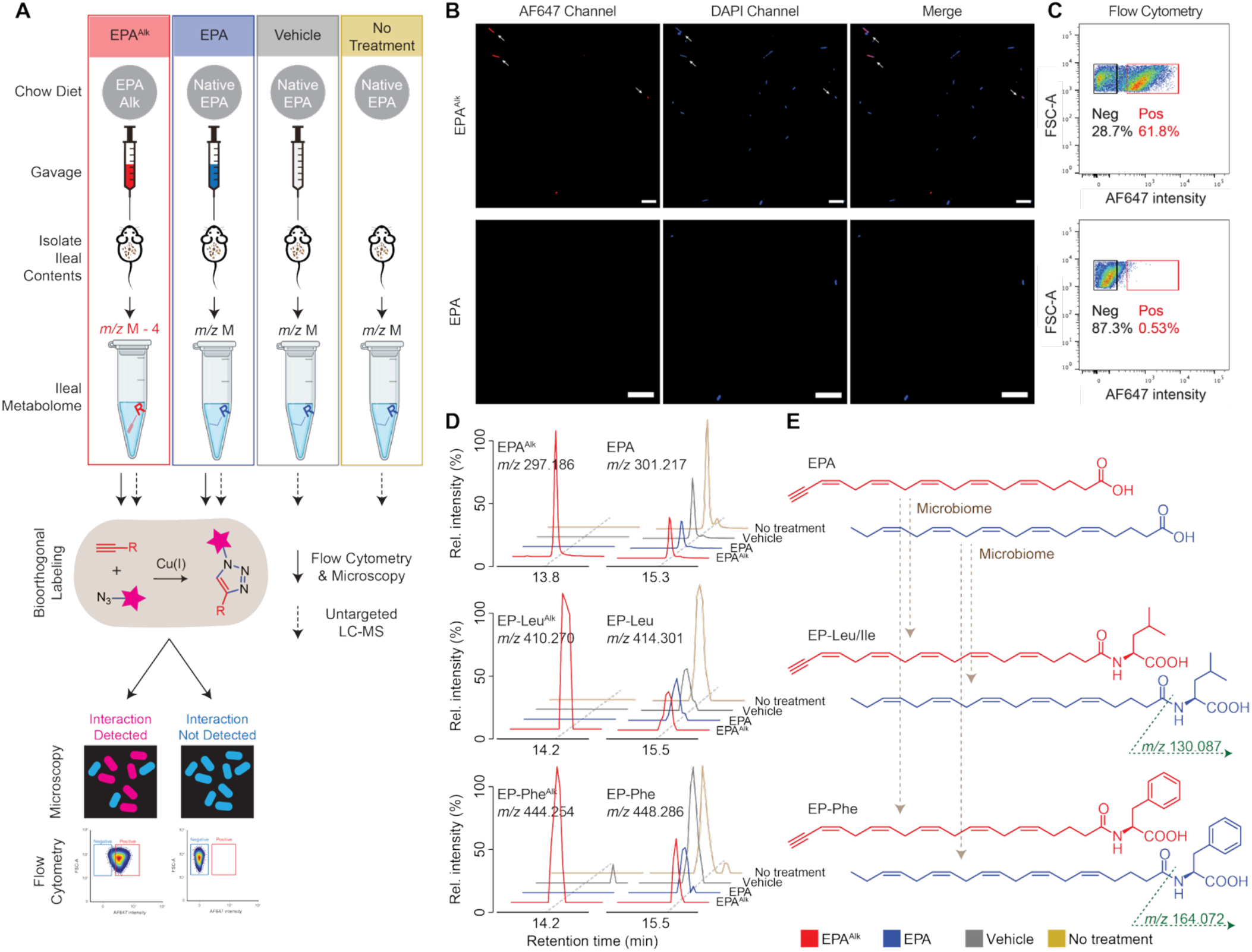
Interaction between ileal gut microbes and EPA detected *in vivo.*

(A) Schematic depicting workflow to detect bacterial EPA^Alk^ interaction *in vivo* via microscopy, flow cytometry, and LC-MS. (B) Fluorescence microscopy images show click-stained bacteria that were isolated from the ileal contents of EPA^Alk^- and native EPA-treated mice. Scale bars in microscope images are 10 μm, and white arrows point to bacterial cells with overlapping blue and red fluorescence. (C) Density plots exhibit flow cytometry analysis of click-stained bacteria isolated from the ileal contents of EPA^Alk^- and native EPA-treated mice. Positive and negative percentages on flow cytometry plots reflect percentages of the cell population that are positive and negative for red fluorescence, respectively. (n = 2 experiments; 2-3 mice per treatment per experiment) (D) EICs depicting the detection of alkyne-modified EP (iso) leucine and EP phenylalanine in EPA^Alk^-treated mouse ileal contents. Native EP (iso)leucine and EP phenylalanine were detected in samples from all treatment groups. (E) Structures of EPA, EP-Leu/Ile and EP-Phe in their native form (blue and the alkyne form (red). Green arrow represents diagnostic fragment detected in the tandem mass spectrometry experiment. Intensities are normalized to the highest intensity value in each chromatogram, which is set to 100%. (n = 5-6 mice per treatment).

As noted for LA, EPA may also undergo microbial conversion to an EPA-derived metabolite. We then further utilized our methodology to identify novel diet-dependent microbial metabolites. The alkyne modification imparts a predetermined mass difference, enabling a streamlined comparative metabolomic platform to reveal EPA- and EPA^Alk^-derivatives. Metabolomes were extracted from the luminal intestinal contents of EPA^Alk^- and native EPA-treated mice to identify features unique to the EPA^Alk^-treated mice. In contrast to the fluorescence-based detection of EPA^Alk^ interactors described above, this comparative metabolomics-based approach leverages the alkyne handle without the need to click Alexa Fluor 647-azide to our metabolites of interest. Mice that received vehicle gavages or no gavages were also included as negative controls for the detection of EPA^Alk^-dependent microbial metabolites. EPA^Alk^ and EPA were detectable by LC-MS in intestinal contents, demonstrating that bacteria residing in the lumen of the distal small intestine encounter dietary EPA (Figure 2D).

Multiple EPA^Alk^-derived metabolic features were detected, confirming that dietary EPA is transformed to other metabolites (Figure 2D). Chemical formulas and fragmentation analysis suggested the attachment of leucine or isoleucine to EPA via an amide bond, creating *N*-eicosapentaenoyl (iso)leucine (EP-Leu) (Figure 2D-E). Another feature suggested the *N*-attachment of phenylalanine to EPA, creating *N*-eicosapentaenoyl phenylalanine (EP-Phe) (Figure 2D-E). As expected, the non-native, alkyne-modified forms of these metabolites were exclusively detected in EPA^Alk^-treated mice (Figure 2D). The native forms of these metabolites were detected in all mice (Figure 2D), likely due to the presence of EPA and EPA precursors in the chow diet. To further confirm the identity of EP-Phe, EP-Leu, and EP-Ile, we performed a straightforward 1-ethyl-3-(3-dimethylaminopropyl)carbodiimide (EDC)-mediated chemical coupling of EPA to Phe, Leu, or Ile. As expected, high-resolution *m/z,* retention times, and fragmentation patterns of EP-Phe, EP-Leu, and EP-Ile generated via chemical synthesis matched those of biological sources from *ex vivo* mouse cultures (Figure S2). From this point forward, EP-Leu and EP-Ile could not be distinguished via LC-MS. Therefore, for the remainder of the text, EP-Leu refers to either EP-Leu or EP-Ile. Past literature describes the production of *N*-acyl amino acids from saturated and monounsaturated fatty acids in soil microbes^32,33^ and gut-associated microbes^34-37^, but, to our knowledge, no previous study has reported on the production of EP amino acids from dietary EPA by gut bacteria.

### EPA uptake in complex microbial communities

To confirm EPA uptake specifically occurring in gut microbes and independent of mammalian host tissues, we used *ex vivo* cultures of mouse stool bacteria to assess bacterial EPA uptake separate from the host. Bacterial cells were isolated from freshly collected mouse stool and were incubated with EPA^Alk^ *ex vivo* for 5 hours (Figure 3A). EPA^Alk^-treated fecal bacteria contained a fraction of bacterial cells positive for EPA uptake (Figure 3B-C). These data indicate that select culturable bacteria from this mouse gut microbial community took up EPA^Alk^ within the 5-hour incubation period. The specialized capacity of certain bacteria to assimilate exogenous EPA reinforces the idea that the presence of LCPUFA-interacting microbes may be an important determinant of the effect of dietary LCPUFA supplementation on gut microbiota activities. Additionally, we confirmed the mixed microbial cultures were capable of EP-Phe and EP-Leu (Figure 3D) production, highlighting that microbial systems convert EPA to EP-Phe/Leu.

**Figure 3.**
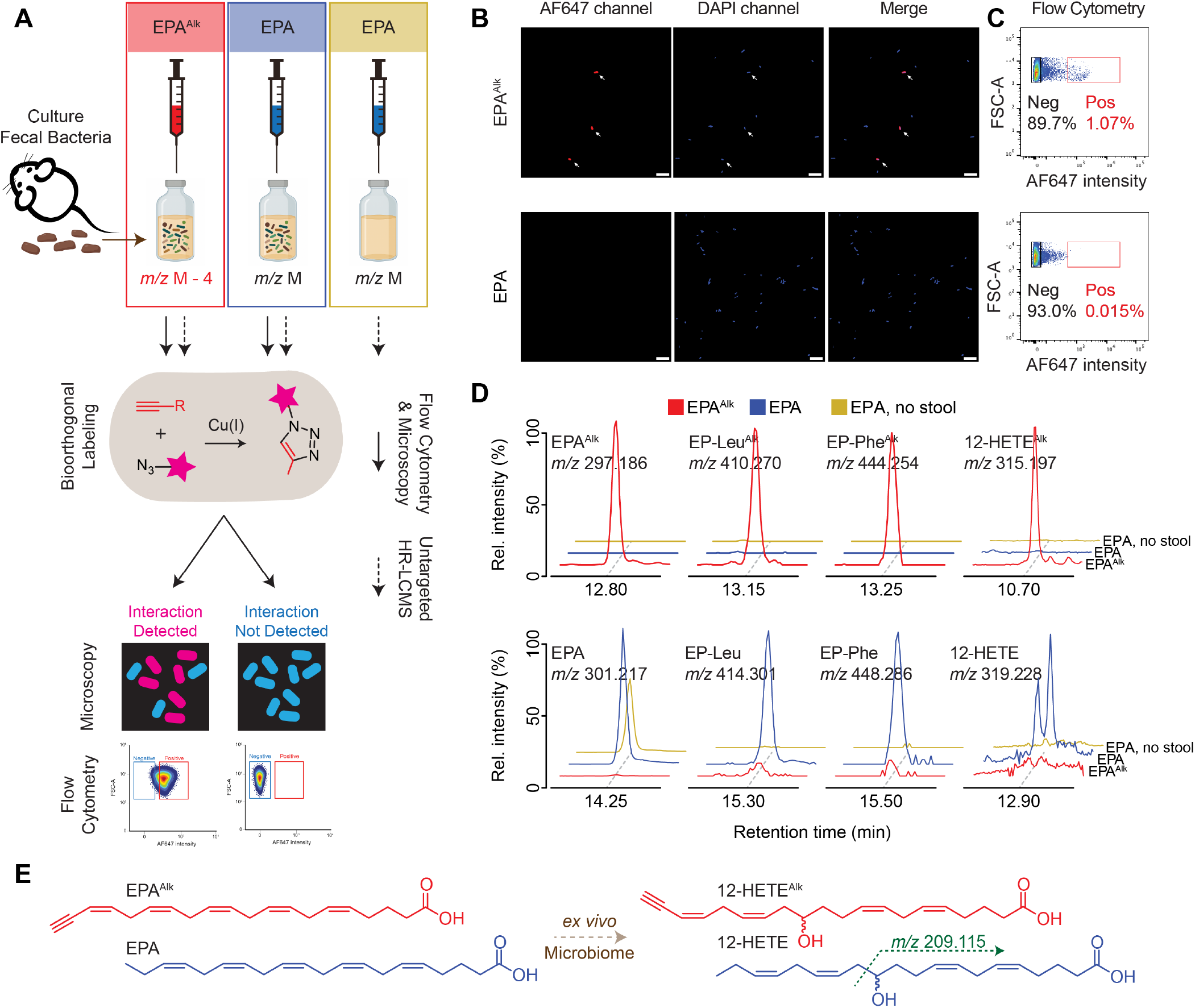
Mouse stool bacteria take up EPA in *ex vivo* cultures.

(A) Schematic depicting workflow to detect bacterial EPA^Alk^ *ex vivo* interactions via microscopy, flow cytometry, and LC-MS. (B) Fluorescence microscopy images show click-stained bacteria from EPA^Alk^-treated or native EPA-treated *ex vivo* cultures. Scale bars in microscope images are 10 μm, and white arrows point to bacterial cells with overlapping blue and red fluorescence. (C) Density plots exhibit flow cytometry analysis of click-stained bacteria isolated from EPA^Alk^-treated or native EPA-treated *ex vivo* cultures. Positive and negative percentages on flow cytometry plots reflect percentages of the cell population that are positive and negative for red fluorescence, respectively. (n = 3 cultures per treatment group) (D) EICs representing the detection of EP-Leu, EP-Phe and a candidate 12-HETE. The alkyne-variants were detected when EPA^Alk^ was available to the system. (E) Structures of EPA, 12-HETE in their native form (blue) and the alkyne form (red). Green arrow represents diagnostic fragment detected in the tandem mass spectrometry experiment.

In addition to metabolites detected in the *in vivo* mouse experiment, we detect hydrated EPA in *ex vivo* cultures (Figure 3D-E). Tandem mass spectrometry suggests hydration of EPA occurs at carbon 12 (Figure 3E). A 12-hydroxy-eicosatetraenoic acid (12-HETE) structure is congruent with the previously reported detection of hydroxy-fatty acids produced by *Lactobacillus* spp^21,22^. Pure standards of 12-hydroxy-5Z,8Z,10E,14Z-eicosatetraenoic acid and 11(12)-epoxy-5Z,8Z,14Z-eicosatrienoic acid were analyzed by LC-MS, and we observed that the bacterial metabolite retention time closely resembled that of a 12-HETE (Figure S3). Past studies describe *Lactobacillus acidophilus* fatty acid hydratase enzymes that convert various LCPUFAs to hydroxy-fatty acids^22,38^. Reactions performed by these enzymes include the conversion of EPA to 12-HETE^38^. Further structural elucidation is needed to ascertain the specific connectivity of the hydrated EPA identified, but this result along with past findings suggest microbes in the gut produce hydroxy-fatty acids from EPA. Identifying *N*-acylated amino acids of EPA and 12-HETE demonstrate the utility of metabolic labeling coupled to comparative metabolomic workflows for accelerating the identification of multiple diet-dependent gut bacterial metabolites.

### Bioorthogonal labeling reveals bacteria that take up EPA and produce EP-Phe/Leu

Given the potential power in identifying microbes that interact with LCPUFAs, as demonstrated by tracing co-occurrences of HYA and HYA-producing bacteria in infant stool (Figure 1D, Figure S1), we sought to survey various bacteria for species-specific EPA interactions. To this end, we elected to screen six gut-associated bacterial strains commonly found in human infants^39-41^, each previously implicated in dietary lipid engagement^24,35,42-47^. Each strain was incubated with EPA^Alk^ for 4 hours in a medium based on MiPro^48^, a medium formulated to sustain a broad range of gut-derived microbes *ex vivo* (Figure 4A). Significant uptake (>20%) of EPA^Alk^ was observed for *Clostridium bifermentans* (Figure 4B), consistent with previous reports that EPA undergoes hydrogenation in pure cultures of *Clostridium bifermentans*^42^. We also note EPA^Alk^ uptake by *Bacteroides thetaiotaomicron* (Figure 4B), a bacterium previously observed to assimilate saturated fatty acids^49^. Increased LCPUFA consumption and serum levels have been correlated with gut *Bacteroides* in rodent models and humans^18,50-52^, but direct interactions between *Bacteroides* species and EPA have not yet been investigated. Another major interactor was *Escherichia coli*, a bacterium previously shown to incorporate exogenous PUFAs into membrane phospholipids^43^. *Bifidobacterium breve, Enterococcus faecalis, Lactobacillus johnsonii* were also determined to be EPA^Alk^ assimilators (Figure 4B, D) but at lower levels than the gram-negative microbes. Due to the potential for membrane-associated EPA to be differentially detected in gram-negative versus gram-positive microbes because of the staining steps used in click reactions for fluorescence-based detection, we were also able to verify uptake purely using LC-MS analysis.

**Figure 4.**
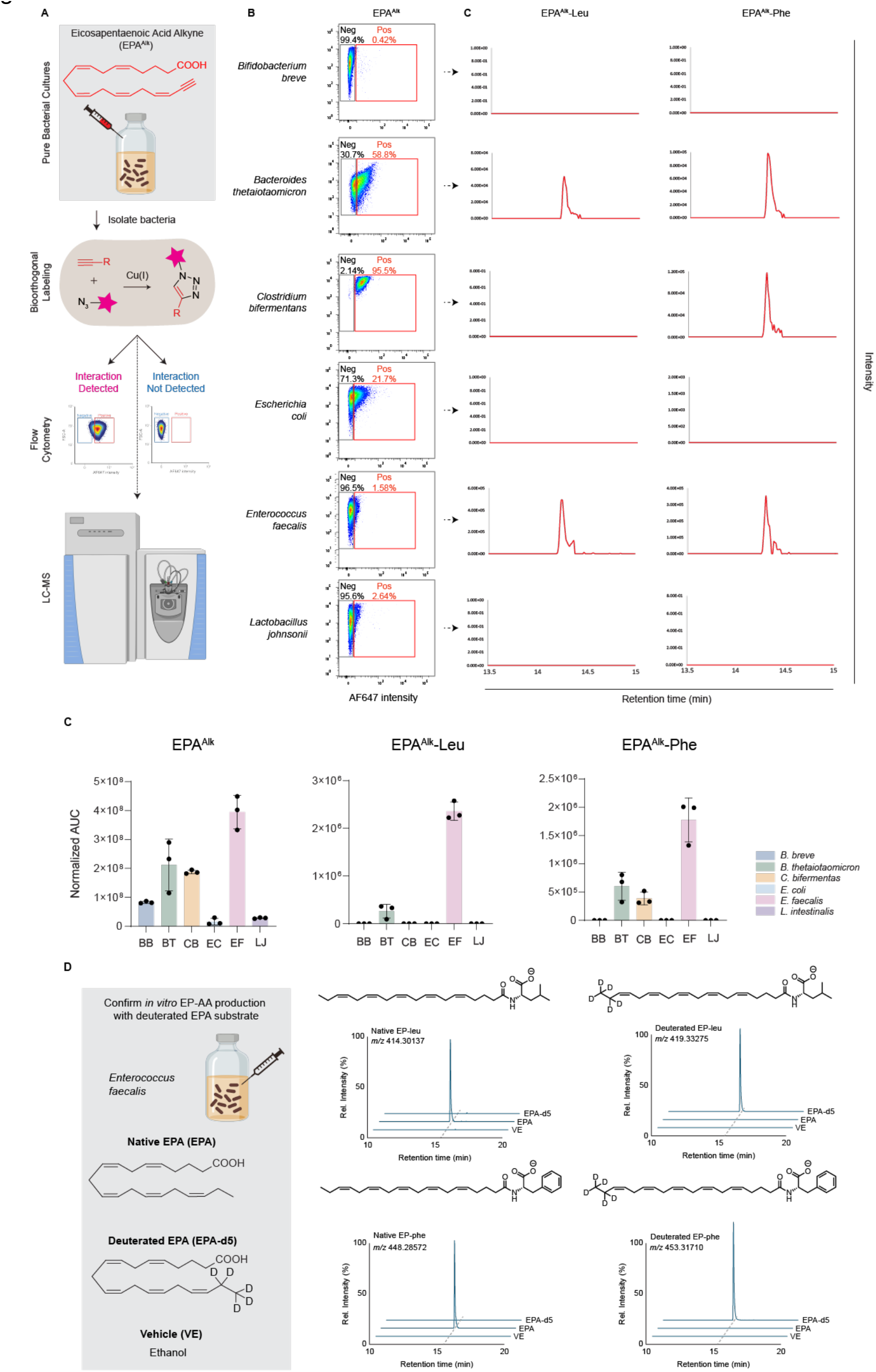
Select gut-associated bacterial species take up exogenous EPA *in vitro*.

(A) Schematic depicting workflow to detect single bacterial species EPA^Alk^ interactions via microscopy and LC-MS. (B) Flow cytometry results of click-staining EPA^Alk^-treated for *B. breve, B. thetaiotaomicron, C. bifermentans, E. coli, E. faecalis,* and *L. johnsonii*. (C) EICs from LC-MS analysis of EPA^Alk^-treated bacterial cultures depicting the detection of EPA^Alk^-Phe or EPA^Alk^-Leu. The intensities are scaled to the highest detected peak. (D) Bar plots of area under curve (AUC) of EPA^Alk^, EPA^Alk^-Phe, and EPA^Alk^-Leu detected by LC-MS in bacterial culture EICs (n = 3 cultures per species). (E) Workflow of using deuterated EPA-d5 to verify the presence of EP amino acids. EICs showed the presence of native and labelled EP amino acids in *in vitro* cultures.

Thus, we then followed up individual species-level characterizations with targeted LC-MS analysis to identify bacteria capable of producing EP amino acids. As expected EPA^Alk^ was detected in all EPA^Alk^-treated bacterial cultures but not in native EPA-treated cultures (Figure 4D). Among the six bacteria surveyed, 3 showed EP-Phe or EP-Leu production, namely *Enterococcus faecalis, Bacteroides thetaiotaomicron,* and *Clostridium bifermentans. Bacteroides thetaiotaomicron, and Clostridium bifermentans* were modest producers of EP-Phe/Leu. *Enterococcus faecalis* was the major producer of EP-Phe/Leu amongst our select panel (Figure 4C,D). We further confirmed the production of EP amino acids by incubating either a native EPA or a deuterated EPA-d5 substrate with *E. faecalis* pure cultures. We were only able to detect the native and the labelled EP-amino acids when the respective substrates were given (Figure 4E). This observation confirms that EP amino acids can be directly converted from EPA by *E. faecalis* in a substrate-dependent manner (Figure 4E). Altogether, these pure isolates reveal a previously undescribed capacity for EPA uptake in gut-associated bacteria and highlight that minor community members may have the potential to exert significant bio-transformation effects on EPA.

### EP amino acids are detected in human infant stool

Enabled with novel gut microbial transformations of EPA, we endeavored to identify EP-Phe and EP-Leu in human infant stool samples. Indeed, EP-Phe and EP-Leu were detected in stool from select infants and at multiple time points within the first 5 months of life (Figure 5A-B, Figure S4). Tandem mass spectrometry and synthetic standards confirmed the identity of EP amino acids in infant stool (Figure S4). EP-Phe and EP-Leu followed similar patterns of detection across infants and were clearly detected in stool from select infants (Figure 5A).

**Figure 5.**
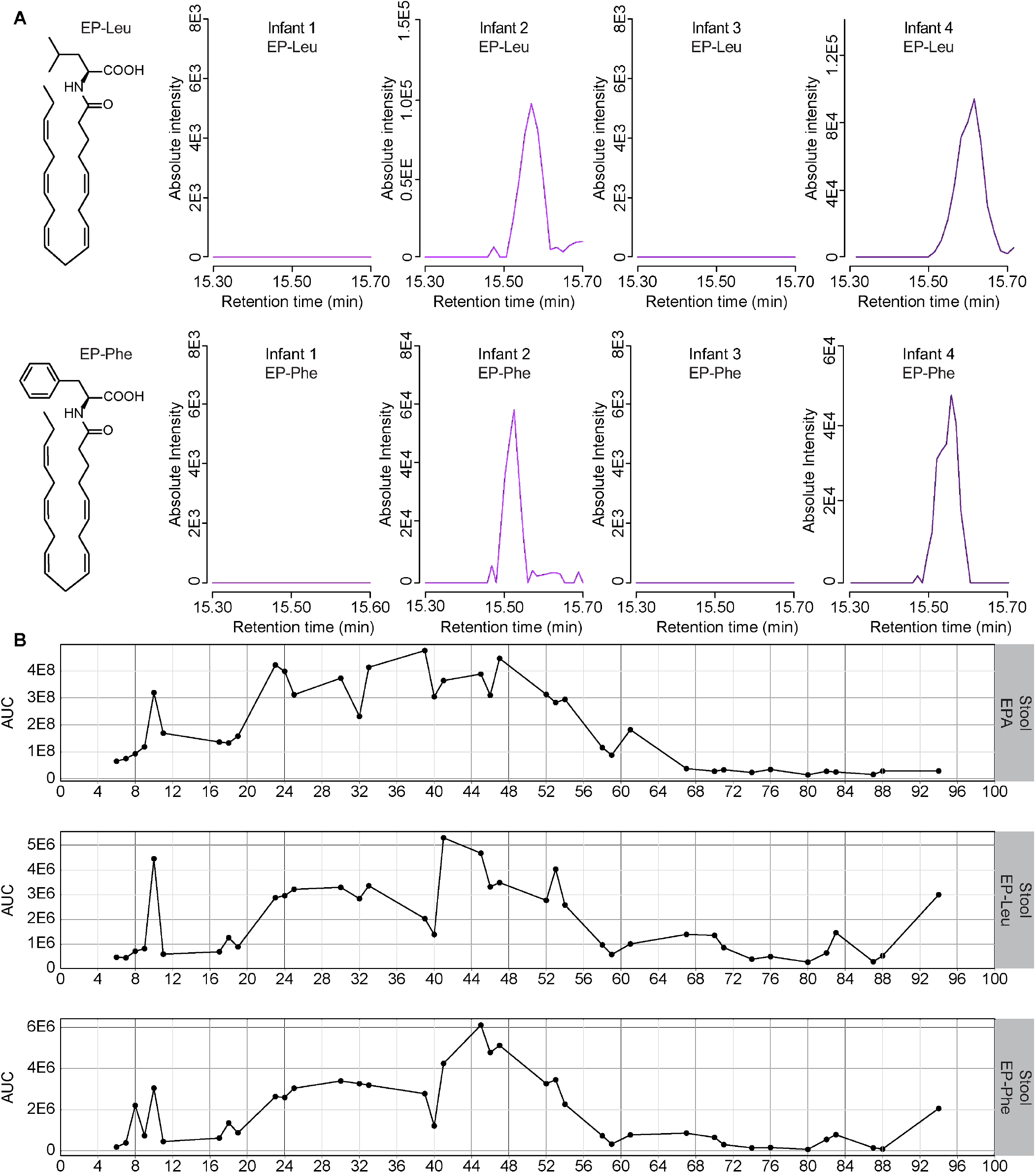
EP amino acids are detected in human infant stool.

(A) EP-Leu and EP-Phe were detected in stool of multiple infants, but EP amino acids were not detected in all infant stool samples. Chromatograms shown are MS1 EICs for the parent ions displayed at the top of each panel. Y-axes display absolute intensities, reflecting peak heights. (B) Levels of EPA, EP-Leu and EP-Phe in Infant 5 stool were assessed at multiple timepoints within the first 100 days of life. Stool EPA, EP-Leu and EP-Phe levels are shown as MS1 peak areas (i.e., area under curve, AUC).

EPA, EP-Leu and EP-Phe abundances in longitudinal stool samples from infant 5 over time were measured due to the exposure of this infant to both human milk and infant formula over time (Figure 5B). EP-Leu and EP-Phe levels in the stool varied in a similar pattern, complementing the similarities in detection noted across different infants (Figure 5B). Tracking the abundance of gut microbial EPA metabolites in infant stool presents the potential to examine how dietary exposure to LCPUFAs may contribute to infant health.

## DISCUSSION

Beyond the hydration and conjugation of LA and ALA described in certain bacterial species, gut microbial metabolism of dietary LCPUFAs has not been extensively characterized. This is particularly true for the omega-3 LCPUFA EPA. In this work we were able to demonstrate that gut bacteria utilize diet-derived EPA *in vivo* using bioorthogonal labeling with EPA^Alk^. Culturing mouse stool bacteria and pure bacterial strains with labeled EPA provided further evidence that gut bacteria directly take up and metabolize EPA independent of host EPA metabolism. Comparative metabolomics of *in vivo, ex vivo,* and pure culture gut bacterial systems showed that gut bacteria produce *N*-acyl amino acids (NAAs) EP-Phe and EP-Leu. Although microbial *N*-acyl amino acids have been previously characterized with fully saturated fatty acids^36^, our work demonstrates the first characterization of *N*-eicosapentaenoyl amino acid production by gut bacteria.

This discovery feeds into the growing roles for NAA in mediating biology in host-microbe systems. This includes one study that revealed NAA production by *Bacteroides thetaiotaomicron* enhanced its ability to colonize the host gut^35^. Another study showed bacterial NAAs act upon certain G-protein-coupled receptors and can thereby influence host physiology^36,37^. Specifically, *N*-3-hydroxypalmitoyl glycine and *N*-palmitoyl alanine are ligands for GPR132^36,37^, which is expressed in various immune cell types and has been implicated in macrophage trafficking^53-55^. The specific investigation of NAA species containing polyunsaturated fatty acyl chains is uniquely important as these lipids have been found to activate GPR132^56^, but the production of these metabolites has not yet been attributed to gut microbes. Furthermore, polyunsaturated fatty acyl amides with structural similarities to the NAAs described in this study have been shown to modulate cell proliferation^57,58^. Relevant to host health, NAAs from the gut microbiome^59^ have been associated with metabolic disease^60^, further solidifying the importance of characterizing of the origin and modifiers of these metabolites.

Given LCPUFA-derived NAAs have been shown to engage in host biology, we envision that gut microbial EP amino acids may also engage host systems. Our study found EP-Phe and EP-Leu were present both in the intestinal contents of mice fed EPA and in stool collected from human infants.

Bioorthogonal labeling of EPA administered to mice confirmed EP-Phe and EP-Leu in the intestinal lumen are produced from EPA consumed orally. Analysis of *ex vivo* stool cultures further validated that gut bacteria are capable of producing EP-Phe and EP-Leu.

In the human infant context, given differences in free EPA concentrations between human milk versus infant formula, gut bacterial EPA uptake may be high in systems where oral PUFAs are less absorbed based on their bioavailability in the upper intestinal tract. Gut bacterial EPA uptake may also be enhanced when infant diets are naturally high or supplemented with EPA. Variability in the EPA content of human milk collected from different individuals has also been reported in other studies^61,62^. Variability in human milk LCPUFA content has been partly attributed to differences in LCPUFA intake in mothers’ diets^62-65^. While previous studies have compared the LCPUFA profile of infant formulas to that of human milk,^29,30^ this study provides more information on these differences by reporting absolute concentrations of free and total LCPUFA contents and by comparing formulas and milk from individual mothers. Measuring LCPUFA levels as a percentage of total fatty acids extracted from milk, past studies on infant formulas suggest similar or lower levels of EPA, DHA and ARA, and similar or higher levels of ALA and LA, compared to human milk^29,30^. Here, measuring LCPUFA levels as milligrams LCPUFA per 100 mL milk, we report comparative absolute concentrations of total EPA, DHA, ARA, and LA between mothers. Both the interindividual variability in human milk LCPUFA content and the differences between human milk and infant formulas raise the question of how these differences impact the metabolic milieu within the infant. Measurements of LCPUFAs in infant stool and labeled EPA in mouse intestinal contents in this study suggest dietary LCPUFAs such as EPA reach distal portions of the intestinal tract where gut microbes reside. This means one consequence of altered forms of LCPUFA consumption could be differences in gut microbial LCPUFA interactions – including EP-Phe and EP-Leu production – which may carry currently unknown implications for infant health and development.

Further characterization of EP amino acid synthesis by gut microbes and the relevance of this activity to human infants will determine whether the presence or absence of this gut microbiome function can directly influence infant health. Given LCPUFAs are being used as therapeutic supplements to improve outcomes in neonatal intensive care units^66^, where infant microbiomes are often altered from the term/healthy infant state^67^, it is important to fully define both the dietary exposure to LCPUFAs and the LCPUFA-responsive microbes that allow for optimal health during infancy. Our investigations into LCPUFA-dependent gut microbial metabolites provide additional functional insight for infant microbiome profiling, fostering a framework for manipulating diet, microbiome composition, and microbial LCPUFA metabolites as therapeutic strategies during infant development with direct relevance to dietary LCPUFA supplements that are also popularly consumed by adults.

## MATERIALS AND METHODS

Materials and methods detailed in supplemental materials

## ACKNOWLEDGEMENTS

This work was funded by a National Institutes of Health (NIH) grant R35GM138281 (E.L.J.) and a Schwartz Fund Visionary Scientist Award (E.L.J). E.L.J. is a Pew Biomedical Scholar. J.M.C.C. was supported by an NIH T32 grant (T32-DK007158). We thank the Genomics Facility of the Biotechnology Resource Center (BRC) of Cornell Institute of Biotechnology for their help with sequencing experiments, and we thank the BRC Proteomics and Metabolomics Facility for their help with select metabolomics experiments. Figures were created, in part, with Biorender.com.

## AUTHOR CONTRIBUTIONS

Conceptualization, J.M.C.C., H.H.L., and E.L.J.; methodology, J.M.C.C., X.T, H.H.L., and E.L.J.; investigation, J.M.C.C., X.T, H.H.L, K.R.B, and S.M.T.; formal analysis, J.M.C.C., X.T, H.H.L., S.M.T.; writing—original draft, J.M.C.C. and E.L.J.; writing—review & editing, J.M.C.C., X.T, H.H.L, and E.L.J.; funding acquisition, E.L.J.; supervision, E.L.J.

## CONFLICTS OF INTEREST

The authors declare no competing interests.

## Notes

### Competing Interest Statement

The authors have declared no competing interest.

