## Supplemental Information for "Dietary omega-3 polyunsaturated fatty acids interact with select gut microbes to produce bioactive metabolites found in infant gut microbiomes"

### MATERIALS AND METHODS

**Chemicals.** Optima HPLC-grade solvents were purchased from Fisher Scientific. The following compounds were purchased from Cayman Chemical to serve as LC-MS standards and as substrates for animal and bacterial culture experiments: eicosapentaenoic acid (EPA), EPA alkyne ($EPA^{Alk}$), EPA-$d_5$, docosahexaenoic acid (DHA), arachidonic acid (ARA), α-linolenic acid (ALA), linoleic acid (LA), (±)-10-hydroxy-12(Z)-octadecenoic acid (HYA), (±)12-hydroxy-5Z,8Z,10E,14Z-eicosatetraenoic acid and *rel*-10-[(2R,3S)-3-(2Z)-2-octen-1-yl-2-oxiranyl]-5Z,8Z-decadienoic acid. For the synthesis of *N*-eicosapentaenoyl (EP) amino acid standards, L-leucine and L-isoleucine were purchased from Sigma-Aldrich, and L-phenylalanine was purchased from Alfa Aesar (Haverhill, Massachusetts).

**Synthesis of EP amino acid standards.** The method used to synthesize EP amino acid standards was based on a previously published method for the derivatization of short chain fatty acids[1]. Solutions of 50 mM *N*-(3-Dimethylaminopropyl)-*N*′-ethylcarbodiimide (EDC) and 25 mM leucine, isoleucine or phenylalanine were prepared in a water-methanol mixture (3:7, v/v). 200 µL 10 mM EPA (in ethanol), 400 µL isopropanol, 300 µL 3.5% pyridine (in methanol), 300 µL 50 mM EDC and 300 µL 25 mM amino acid were combined (1.5 mL total volume). Reactions were incubated at 37°C for 30 min. Reaction solutions were then dried completely in a Thermo Scientific Savant SpeedVac vacuum concentrator. Dried reaction products were reconstituted in 200 µL methanol. Solutions were centrifuged at 18213 x g at 4°C for 30 min to remove precipitate, and 150 µL supernatant was collected. Methanol solutions were dried in a SpeedVac and reconstituted in acetonitrile. Partial fractionation of reaction mixes was performed by passing mixes through a silica column containing 12 g silica gel powder and acetonitrile. Fractions were dried completely in a SpeedVac and concentrated in 200 µL methanol. Fractions were analyzed on a Thermo Scientific Vanquish Horizon UHPLC System coupled with a Thermo Scientific TSQ Quantis Triple Quadrupole mass spectrometer (MS). For detection of EP amino acids in fractions, the following gradient of 0.1% formic acid in water (solvent A) and 0.1% formic acid in acetonitrile (solvent B) at a flow rate of 0.4 mL/min was applied to an InfinityLab Poroshell 120 EC-C18 column (50 mm x 2.1 mm, 2.7 µm, Agilent, part number 699775-902) maintained at 50°C: 35% B for 1 min, linear increase to 100% B at 19 min, hold at 100% B for 5.9 min, linear decrease to 20% B at 25 min and hold at 20% B until 27 min. Mass spectrometer parameters were as follows: full scan; negative mode; HESI source; spray voltage of 3.5 kV; ion transfer tube temperature at 300°C; vaporizer temperature at 350°C; sheath, aux, and sweep gases set to 50, 10, and 1, respectively; scan rate of 1000 Da/sec; and resolution 0.7. The m/z scan range was 200 to 1000. Fractions where EP amino acid was present were pooled, dried completely in a SpeedVac and resuspended in 200 µL methanol. Final fractionation was performed using the Vanquish UHPLC coupled with a Thermo Scientific Fraction Collector (Model ASX-280).



^Current Affiliation: Salk Institute for Biological Sciences, San Diego, CA

The EP amino acid solution was injected, 24 µL at a time, and run through a Hypersil GOLD C18 preparative HPLC column (250 mm x 10 mm, 5 µm, Thermo Scientific, part number 25005-259070A) maintained at 50°C using the same mobile phases described above. The following gradient was used for fraction collection: 50% B at flow rate 2 mL/min for 2 min, linear increase to 100% B at flow rate 4 mL/min at 25 min, hold at 100% B at flow rate 4 mL/min for 23.9 min, linear decrease to 50% B at flow rate 2 mL/min at 49 min and hold at 50% B at flow rate 2 mL/min until 50 min. Fractions containing EP amino acid were pooled, dried completely in a SpeedVac and concentrated in 200 µL ethanol. The presence of EP amino acid was verified using the same LC-MS full scan method used to analyze fractions from partial fractionation. Tandem mass spectrometry (with collision energy set to 22 V) was also used to confirm EP amino acid presence and identities. Purified EP amino acid was stored at -80°C for long-term use.

**Animal experiments.** All mouse experiments were performed according to a protocol approved by the Cornell University Institutional Animal Care and Use Committee (protocol no. 2010-0065).

**Culturing mouse stool bacteria *ex vivo*.** Fresh mouse fecal pellets were immediately transferred to sterile pre-reduced PBS containing 0.1% L-cysteine hydrochloride, pre-warmed to 37°C. Fecal pellets were collected at a concentration of approximately 1 g stool per 15 mL 0.1% L-cysteine. Stool was suspended in 0.1% L-cysteine with intermittent brief vortexing for a total of 30 sec. The stool suspension was transferred to an anaerobic chamber containing 70% $N_2$, 25% $CO_2$ and 5% $H_2$. The stool suspension was allowed to sit for 3 min to allow solid debris to settle, and the fecal slurry above the debris was transferred to a new sterile tube. Bottles containing 25 mL pre-warmed anaerobic Modified Tween Medium (MTM) broth were inoculated with 500 µL fecal slurry. MTM consisted of the following: 10 g/L yeast extract, 400 mg/L potassium phosphate dibasic, 400 mg/L potassium phosphate monobasic, 0.1 mg/L magnesium sulfate heptahydrate, 0.008 mg/L calcium chloride dihydrate, 0.025% Tween 80, 1 mg/L hemin, 500 mg/L L-cysteine hydrochloride and 0.001% vitamin K1. MTM was also supplemented with filter-sterilized 0.2% casamino acid before bacterial inoculation. Fecal bacteria were grown for 12h in MTM at 37°C before EPA or EPAAlk administration. 10µM EPA or $EPA^{Alk}$ was administered for fluorescence labeling experiments, while 1µM EPA or $EPA^{Alk}$ was administered for untargeted metabolomics experiments. Cultures were incubated with treatments for 5 h. Each treatment condition was performed in triplicate.

**Bacterial cultures incubated with $EPA^{Alk}$.** Pure bacterial strains were stored in frozen glycerol stocks at -80°C. *Bacteroides thetaiotaomicron* (BT), *Escherichia coli* (EC), and *Enterococcus faecalis* (EF) were inoculated and maintained in sterile anaerobic BHIS medium (i.e., Brain Heart Infusion broth supplemented with 5 g/L yeast extract, 10 mg/L hemin, 1 mg/L resazurin, 500 mg/L L-cysteine hydrochloride and 1 mg/L vitamin K3). *Lactobacillus johnsonii* (LJ) was maintained in MRS broth, and *Bifidobacterium breve* (BB) and *Clostridium bifermentans* (CB) were maintained in Reinforced Clostridial broth (ATCC Medium No. 1053), all in anaerobic conditions. After growing bacterial strains overnight in the aforementioned media, 0.5 mL overnight culture was used to inoculate 20 mL anaerobic MTM broth. All strains were allowed to grow for 15 h at 37°C before either EPA or $EPA^{Alk}$ treatment was administered. 10 µM EPA or $EPA^{Alk}$ was administered for fluorescence labeling and comparative metabolomics experiments. Each bacterial strain and treatment combination was performed with 3 replicates. Pure bacterial cultures were incubated with treatments for 4 hours. Following treatment incubation, cultures were collected and processed for click staining or metabolomics, as described below.

**Administration of $EPA^{Alk}$ to mice.** Female, conventionally raised, murine pathogen-free Swiss Webster mice were purchased from Taconic Biosciences at 5 weeks of age and were acclimated for 4 days before performing experiments. Mice were housed in a climate-controlled environment with 12-h light and dark cycles and had ad libitum access to autoclaved standard breeder diet (LabDiet 5021) and to autoclaved water. 8 of the 21 total mice were weighed to estimate the average mouse weight and treatment dose per kg body weight. Mice were housed 3 or 5 per cage and were randomly assigned to receive one of four treatments daily for 3 consecutive days (with 5 or 6 mice per treatment group): no gavage, vehicle (ethanol-PBS, 50:50, v/v) oral gavage, or 160 µmol per kg body weight of either EPA or $EPA^{Alk}$ administered via oral gavage. Each gavage was 100 µL in volume administered using a 20G gavage needle (Fine Science Tools), and all fatty acids were suspended in ethanol-PBS (50:50, v/v) for gavages. On the day of the final gavage, chow was removed 1 h prior to gavages. Mice were fasted for 5 h before euthanasia via decapitation (with euthanasia taking place 4 h post-gavage).

**Collection of mouse tissues.** Following euthanasia, small intestines were divided into thirds, and the most distal section was collected to represent the ileum. The cecum was also collected separately. Luminal contents were gently extruded from intestinal tissues using forceps. Intestinal contents were divided between two tubes (an empty tube for click staining and a pre-weighed tube with 1-mm zirconium beads for metabolome extraction), were snap-frozen in liquid nitrogen and were stored at -80°C.

**Copper(I)-catalyzed azide-alkyne cycloaddition reaction staining (i.e., click reaction staining).** Pure bacterial strain cultures and *ex vivo* stool cultures were centrifuged in 50-mL conical tubes at 3220 x g for 20 min or at 8000 x g for 5 min at room temperature to pellet bacterial cells, and media supernatants were removed. Pre-reduced PBS containing 0.1% L-cysteine hydrochloride was used to resuspend and transfer bacterial cells to autoclaved 1.5-mL centrifuge tubes. Mouse intestinal contents were suspended in 0.1% L-cysteine with intermittent brief vortexing for a total of 20 sec and with sonication at an amplitude of 1 for 10 sec total (pulse on for 1 sec, pulse off for 2 sec). Suspended mouse intestinal contents were centrifuged at 200 x g for 30 sec to separate the bacterial cell suspension from solid debris. The supernatant containing bacterial cells was transferred to a new tube. Bacterial cells were pelleted by centrifuging at 8000 x g for 5 min. Washing and centrifuging of intestinal contents was repeated 2 more times to maximize bacterial cell recovery. Following bacterial cell isolation for both cultures and mouse samples, cells were fixed in 1 mL 10% formalin for 10 min. Formalin was removed by centrifuging bacterial cell suspensions at 8000 x g for 5 min and discarding the supernatant. Bacterial cells were resuspended and washed with filter-sterilized 1% bovine serum albumin (BSA) in PBS 4 times, using the same centrifugation parameters above (i.e., 8000 x g for 5 min at room temperature) to remove washes. Bacterial cells were stained with 5 µM Alexa Fluor 647 (AF647) azide (Invitrogen) using a Click-&-Go Cell Reaction Buffer Kit (Vector Laboratories) according to manufacturer's instructions. Staining solution was removed, and cells were washed six more times with 1% BSA in PBS, all using the same centrifugation parameters above (i.e., 8000 x g for 5 min at room temperature). The final cell pellet was resuspended in sterile PBS containing 0.1% BSA and 2 mM EDTA, and samples were stored at 4°C until microscopy and flow cytometry were performed.

**Flow cytometry analysis.** Click-stained bacterial cells suspended in PBS containing 0.1% BSA and 2 mM EDTA were passed through the 35-µm nylon mesh cap on sterile FACS tubes (Falcon). Filtered cell suspensions were stored on ice, protected from light, when not in use. The optical density (OD) at wavelength 600 nm of filtered cell suspensions was measured to approximate the colony-forming unit (CFU) concentration, according to a standard curve of CFUs versus OD created using pure cultures of *Bacteroides thetaiotaomicron*. Cell suspensions were diluted to achieve a concentration of approximately 5e+06 CFU/mL for flow cytometry analysis on a FACSMelody Cell Sorter (BD Biosciences). The FACSMelody was sterilized by running 70% ethanol through the flow cell, and the instrument was kept under sterile conditions during sorting using sterile PBS as the sheath fluid. Sterile buffer (i.e., PBS containing 0.1% BSA and 2 mM EDTA) along with Spherotech Nano Polystyrene Size Standard Kit 0.58- and 1.32-µm beads (diluted in buffer) were used to set the minimum side scatter height (SSC-H) threshold for event inclusion, allowing for inclusion of bacterial cell-sized events while limiting background noise. Single bacterial cells were selected for using FACS gates on SSC and FSC plots. AF647 dye was excited using a 640-nm red laser, and fluorescence was captured with a 660-nm/20-nm filter. Fixed $EPA^{Alk}$-treated *Bacteroides thetaiotaomicron* and *Lactobacillus salivarius* cells were click-stained alongside *ex vivo* culture and mouse intestinal content samples to serve as positive and negative controls for AF647 fluorescence, respectively, given fluorescence was consistently detected for *B. thetaiotaomicron* and not detected for *L. salivarius* following incubation with $EPA^{Alk}$. Unstained bacterial cells along with stained bacterial cells from EPA-treated conditions served as additional negative controls for AF647 fluorescence, helping to define negative FACS gates and minimize false positive events in positive FACS gates (limited to less than 1% in negative control samples). Flow cytometry data were analyzed using FlowJo software (v.10.10.0).

**Microscopy.** Click-stained bacterial cells suspended in PBS containing 0.1% BSA and 2 mM EDTA were pelleted by centrifugation at 8000 x g for 5 min at room temperature. The supernatant was discarded, and cells were resuspended and diluted in sterile PBS. 5 µL cell suspension was spread evenly on a microscope slide. This was allowed to dry for 30 min protected from light. 5 µL SlowFade™ Diamond Antifade Mountant with DAPI was pipetted onto the slide, and this was covered with a cover slip. Alternatively, cells were stained with Hoechst 33342 trihydrochloride trihydrate: cell pellets were suspended in 1 mL 10 µg/mL Hoechst 33342, were incubated with Hoechst stain for 10 min (rocking and protected from light), were washed once with sterile 1% BSA in PBS, and then were diluted in sterile PBS to be loaded on a microscope slide. Slides were imaged on a Leica DM6 upright microscope.

**Collection of stool and milk from human subjects.** Human stool samples analyzed for the presence of microbial LCPUFA metabolites came from a study measuring stool metabolites during the first 5 months of life. The study protocol was approved by the Cornell Institutional Review Board for Human Subjects Research (protocol no. 2007009697), and all participants provided written informed consent. For infant participants, consent was provided by their immediate caregiver. Stool samples were collected from the diapers of infants 0-5 months of age by their caregivers. Stool samples were collected from the diaper using a sterile stool collection tube containing a top with a scooping spoon (Sarstedt). Participants were instructed to collect human milk samples from the first feed of the day through full expression of milk from one breast and partitioning ~15 mL of that expression into a milk storage bag. Stool and human milk samples were stored in participants' freezers until being shipped to the lab on dry ice. Human milk and stool samples were then stored at -80°C until DNA and metabolomes were extracted.

**Shotgun metagenomic sequencing of infant stool.** DNA was extracted from infant stool using the QIAGEN DNeasy PowerSoil Pro Kit. DNA concentrations were then measured using the Qubit 1X dsDNA High Sensitivity Assay Kit from Invitrogen, and DNA samples were diluted to 0.2 ng/µl to prepare shotgun metagenomic libraries using the Nextera XT DNA Library Preparation Kit from Illumina. The quality and size of DNA libraries were assessed using an Agilent 5200 Fragment Analyzer system. Library DNA concentrations were quantified via Qubit assay, and libraries were pooled for single end 75 bp sequencing on an Illumina NextSeq 500 System. KneadData (v.0.11.0) was applied to raw FASTQ files to trim reads using Trimmomatic and filter out human DNA sequences using bowtie2 (v.2.4.5). (https://github.com/biobakery/kneaddata) For trimming, the minimum average sequence quality score (Phred) was set to 25. MetaPhlAn (v.4.0)[2] (with standard settings) was used to assign taxonomic classifications to processed reads (i.e., KneadData output with repeats removed) and to calculate taxon relative abundances.

**Metabolome extractions.** Mouse intestinal contents were lyophilized in tubes containing 1-mm zirconium beads. Sample weights were measured to determine dry weights for normalization. Methanol was added to each tube for metabolome extraction. Samples were bead beaten in 30-sec intervals for a total of 2 min, incubating samples on ice between bead beatings to prevent overheating. Samples were then sonicated for 3 min at an amplitude of 100 (pulse on for 3 sec, pulse off for 2 sec) and centrifuged at 18,213 x g for 30 min at 4°C. Supernatants were transferred to new tubes and dried completely in a SpeedVac. Dried metabolite extracts were reconstituted in 100 µl methanol, sonicated for 3 min, and centrifuged under the same conditions described above. The resulting supernatants were transferred to LC-MS vials for analysis. For metabolite extraction from bacterial cultures, bacterial pellets were snap-frozen in liquid nitrogen and lyophilized. Metabolites were then extracted using the same procedure described above, and samples were normalized based on protein concentration. For metabolite extraction from infant stool samples, stools were lyophilized and weighed for dry-mass normalization. Metabolites were extracted in methanol, except for the infant stool samples shown in Fig. 5, which were extracted in acetone using the same procedure. Samples were sonicated and centrifuged as described above. The resulting extracts were dried completely using a SpeedVac, reconstituted in 100 µl methanol, sonicated for 3 min, and centrifuged under the same conditions. Supernatants were transferred to LC-MS vials for analysis.

For LCPUFA extraction from milk, first free fatty acids and complex lipids were extracted by combining 100 µL milk with 4 mL chloroform-methanol mixture (1:2, v/v) in glass tubes. Infant formulas were prepared according to manufacturers' instructions. The chloroform-methanol mixture contained enough EPA-$d_5$ to spike each sample with 40 pmol EPA-$d_5$ for internal standard normalization. Samples were vortexed, mixed on a rocking platform for 2 h and then centrifuged at 1500 x g at 4°C for 15 min. Supernatants were transferred to new glass tubes and dried completely using a nitrogen evaporator. Complex lipids were then hydrolyzed using a previously described method.[3] Briefly, 600 µL 1.75 M KOH in ethanol was added to each lipid extract, and samples were incubated at 70°C for 2 h to saponify fats. 400 µL water and 200 µL hexanes were added, and samples were centrifuged briefly to extract saponified fats in the bottom aqueous alcohol layer. 6 N HCl was added to the aqueous alcohol extract to achieve a pH of 1, protonating fatty acids. 600 µL hexanes were added, and samples were vortexed and centrifuged briefly to extract free fatty acids in the upper organic layer. The fatty acid extract was washed with 1 mL water, dried completely in a nitrogen evaporator and concentrated in 200 µL methanol. Samples were centrifuged at 18213 x g at 4°C for 30 min before being transferred to HPLC vials. Samples were stored at -80°C until analyses were performed.

Following metabolomic analysis and peak integration in Skyline, MS1 peak areas of LCPUFAs were normalized to the peak area of the internal standard (EPA-$d_5$). Normalized peak areas were subsequently converted to concentrations using individual standard curves prepared from five serial concentrations of each LCPUFA standard. Bar graphs of LCPUFA concentrations for each sample were then generated in GraphPad Prism 10.

**Metabolomics analyses.** All metabolomics samples except for infant 5 stool samples were run on a Thermo Scientific Vanquish Horizon UHPLC System coupled with either a Thermo Scientific Q Exactive HF Orbitrap mass spectrometer or a Q Exactive Orbitrap mass spectrometer. Infant 5 stool samples were run on a Thermo Scientific UltiMate 3000 HPLC System coupled with a Q Exactive Orbitrap. 3 µL metabolome extract was injected for MS1 analyses, and 8 µL extract was injected for tandem mass spectrometry analyses. For liquid chromatography, a Kinetex EVO C18 column (150 mm x 2.1 mm, 1.7 µm, Phenomenex, part number 00F-4726-AN) maintained at 40°C was used. Solvent A was 0.1% formic acid in water, and solvent B was 0.1% formic acid in acetonitrile. For analyses performed on Vanquish UHPLC Systems, the A/B gradient at a flow rate of 0.5 mL/min started at 10% B for 2 min, increased linearly to 100% B at 20 min, was held at 100% B for 8 min, decreased linearly to 10% B at 28.1 min and was held at 10% B until 32 min. For analyses performed on the UltiMate 3000 HPLC, the A/B gradient at a flow rate of 0.3 mL/min started at 10% B for 2 min, increased linearly to 100% B at 32 min, was held at 100% B for 16 min, decreased linearly to 10% B at 48.1 min and was held at 10% B until 50 min. Mass spectrometer parameters were as follows for MS1 analyses: negative mode; HESI source; spray voltage of 3.0 kV; capillary temperature at

380°C; prober heater temperature at 400°C; sheath, auxiliary, and spare gases set to 60, 20, and 1, respectively; S-lens RF level at 50; resolution 140,000 at m/z 200; AGC target of 3e+06; and max ion injection time of 200 ms. The m/z scan range was 100 to 1000. Tandem mass spectrometry was carried out with an inclusion list of select features using a PRM method that had the same parameters as those described above with the following additions or adjustments: resolution 60,000 at m/z 200; AGC target of 5e+05; max ion injection time of 80 ms; isolation window of 0.7 m/z; stepped normalized collision energies at 25, 35 and 45. Q Exactive instruments were calibrated with calibration solutions from Thermo Fisher Scientific. Raw data files were converted to MZML files using the ProteoWizard MSConvert GUI.[4] For untargeted metabolomics analysis of metabolites in bacterial cultures, peak detection and integration was performed on MZML files using MZmine (v.4.1.0).[5] The aligned feature list exported from MZmine was imported into R, where features were filtered to identify potential EPA- and $EPA^{Alk}$-derived metabolites (Table S1). For targeted metabolomics analyses, peak integration was performed in Skyline.[6]

### SUPPLEMENTAL FIGURES

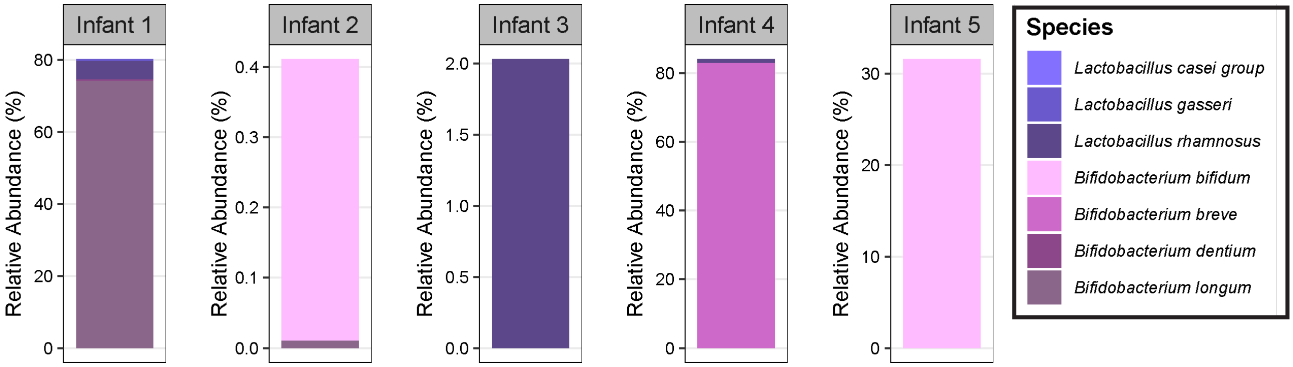


**Figure S1. | Putative HYA-producing bacteria detected in infant stool samples in which HYA was also detected.** Subset of 5 infant stool samples in which HYA was detected were also subjected to shotgun metagenomic sequencing to identify bacterial species present in the infant gut microbiome. *Lactobacillus* and *Bifidobacterium*, genera that include species reported to produce HYA, were detected in these stool samples.

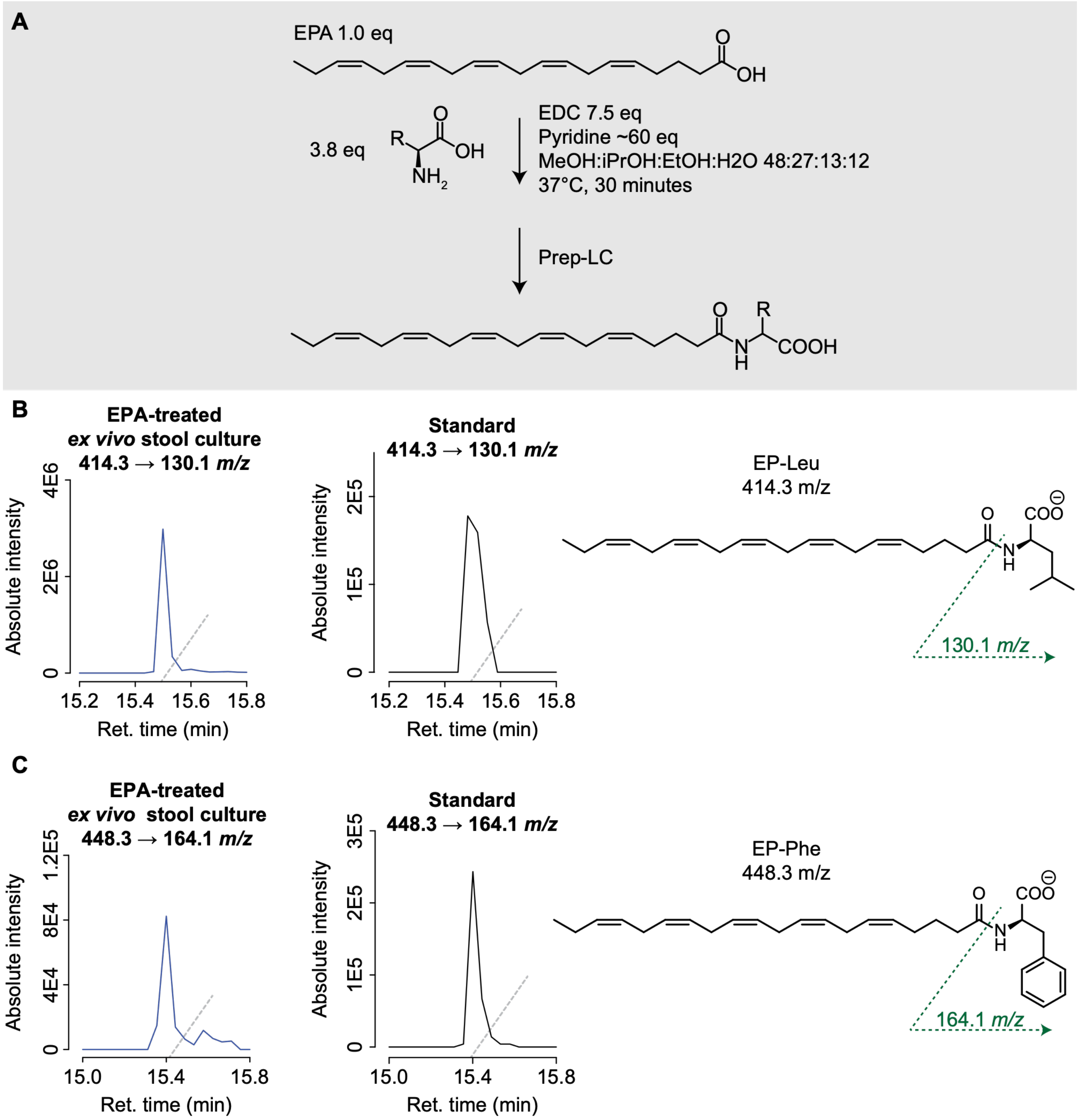


**Figure S2. | Chemical synthesis and LC-MS/MS validation of stool derived EP-Leu and EP-Phe.** (A) Reference standards were synthesized by EDC-mediated chemical coupling and purified by preparative LC. (B) LC-MS/MS chromatograms of EP-Leu from the EPA-treated ex vivo stool culture and the synthesized standards using the transition of *m/z* 414.3 → 130.1 to show matching retention times and product-ion formation. (C) Corresponding LC-MS/MS analysis of EP-Phe using the transition m/z 448.4 → 164.1 which shows matching RT and ion-products of the synthesized standard compared to the EPA-treated *ex vivo* stool culture. MS2 EICs are shown for the fragment ions displayed. Extracted ion chromatograms are filtered to only show the signals associated with the corresponding parent ion. Y-axes display absolute intensities, reflecting peak heights.

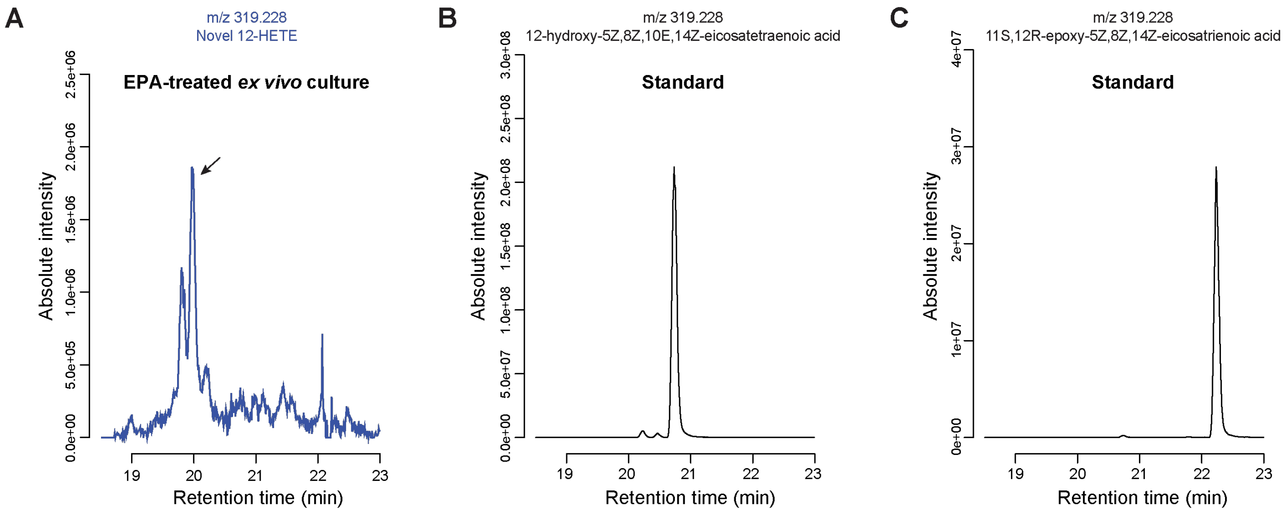


**Figure S3. | Retention time of novel 12-HETE metabolite approaches retention time of 12-hydroxy-5Z,8Z,10E,14Z-eicosatetraenoic acid standard.** An EPA-treated *ex vivo* stool culture sample (A) was run on an LC-MS system alongside commercial standards for 12-hydroxy-5Z,8Z,10E,14Z-eicosatetraenoic acid (B) and 11(12)-epoxy-5Z,8Z,14Z-eicosatrienoic acid (C) using a 50-min method. The retention time of the novel 12-HETE metabolite detected in stool cultures was 20.0 min, while retention times for 12-hydroxy-5Z,8Z,10E,14Z-eicosatetraenoic acid and 11(12)-epoxy-5Z,8Z,14Z-eicosatrienoic acid were 20.7 and 22.2 min, respectively.

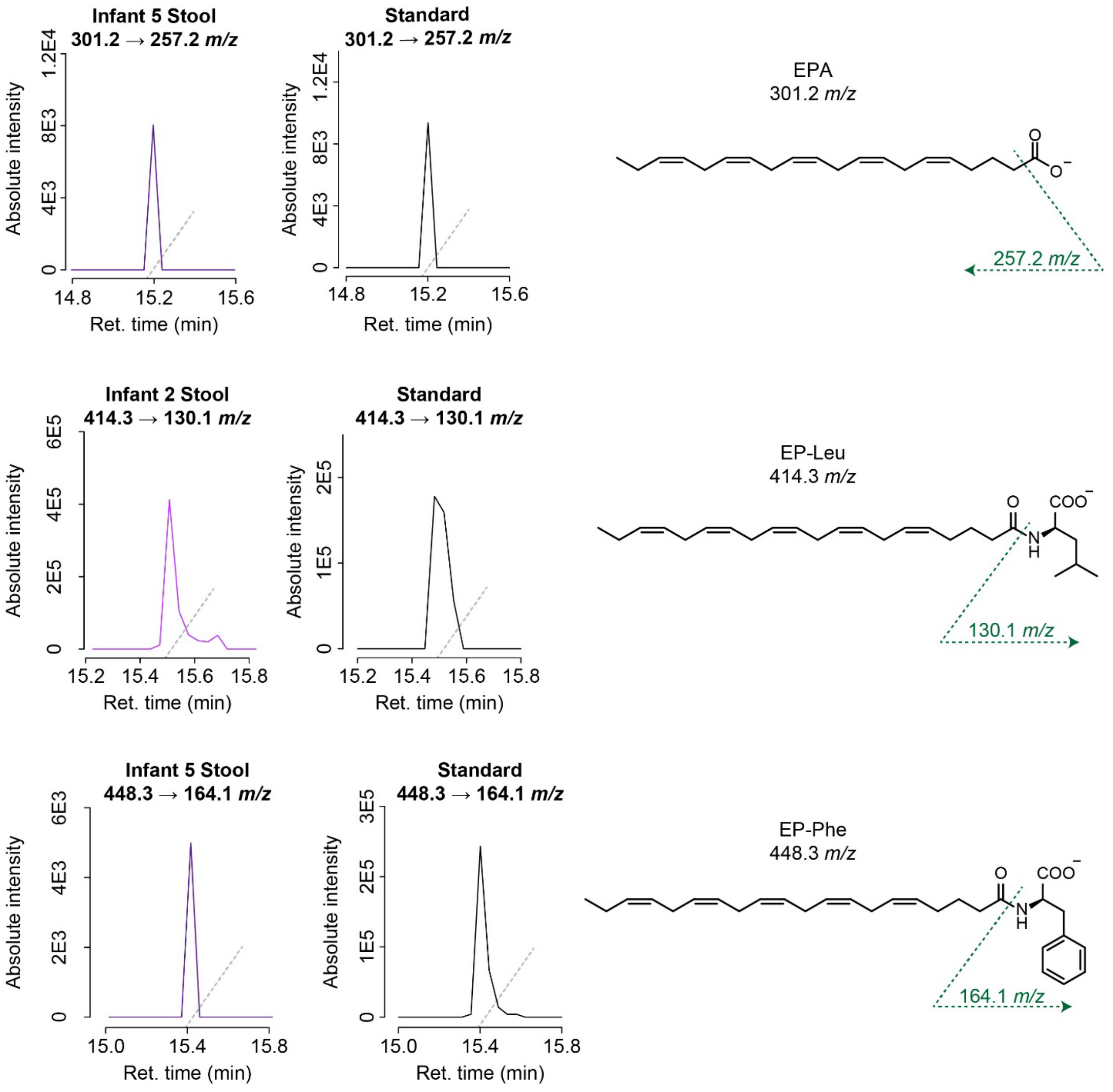


**Figure S4. | Identity of EPA, EP-Leu and EP-Phe peaks confirmed with tandem mass spectrometry and standards.** The same fragment ions associated with EPA (A), EP-leu (B) and EP-phe (C) in *ex vivo* mouse stool cultures (Figure S2) and in standard solutions were detected in infant stool samples, confirming the identity of EPA, EP-leu and EP-phe peaks detected in infant stool. MS2 chromatograms shown are EICs for the fragment ions displayed. EICs are filtered to only show the signals associated with the corresponding parent ion. Y-axes display absolute intensities, reflecting peak heights.

| **Filtering metabolomics features for identification of EPA$^{Alk}$- and EPA-derived metabolites in *ex vivo* mouse stool cultures** | |
|---|---|
| **Filtering criteria (applied in order listed)** | **Number of features passing filter** |
| None | 30,271 |
| Features not detected in solvent blank | 29,302 |

| | |
|---|---|
| Features detected in all 3 $EPA^{Alk}$-treated cultures AND not detected in 2 or more EPA-treated cultures AND not detected in 2 or more EPA-treated sterile medium samples (i.e., candidate $EPA^{Alk}$ derivatives) | 539 |
| Candidate $EPA^{Alk}$ derivatives with corresponding candidate EPA derivatives that meet the following criteria:<br>• detected in all 3 EPA-treated cultures<br>• [not detected in 2 or more $EPA^{Alk}$-treated cultures OR detected in higher abundance in EPA-treated cultures than in $EPA^{Alk}$-treated cultures ($p \leq 0.05$)] AND [not detected in 2 or more EPA-treated sterile medium samples OR detected in higher abundance in EPA-treated cultures than in EPA-treated sterile media ($p \leq 0.05$)]<br>• m/z 4.0313 (+/- 5 ppm) mass units greater than m/z of $EPA^{Alk}$ derivative<br>• elutes before EPA if $EPA^{Alk}$ derivative elutes before $EPA^{Alk}$ OR elutes after EPA if $EPA^{Alk}$ derivative elutes after $EPA^{Alk}$ | 17 |
| Candidate $EPA^{Alk}$ derivatives that do not coelute with $EPA^{Alk}$ AND have EPA equivalents that do not coelute with EPA (since features coeluting with $EPA^{Alk}$ and EPA could be in-source fragmentation artifacts) | 13 |

**Table S1. | Filtering metabolomics features for identification of $EPA^{Alk}$- and EPA-derived metabolites in *ex vivo* mouse stool cultures**. Not detected is defined as either MZmine feature state being "UNKNOWN" or MZmine feature state being "ESTIMATED" and MS1 EIC peak area being less than 1e+04. Detected is defined as either MZmine feature state being "DETECTED" or MZmine feature state being "ESTIMATED" and MS1 EIC peak area being greater than or equal to 1e+04.

Unifying Software Package for Quantitative Metabolomics. J Proteome Res *19*, 1447–1458. 10.1021/acs.jproteome.9b00640.
